# OSpRad; an open-source, low-cost, high-sensitivity spectroradiometer

**DOI:** 10.1101/2022.12.09.519768

**Authors:** Jolyon Troscianko

## Abstract

Spectroradiometery is a vital tool in a wide range of biological, physical, astronomical and medical fields, yet its cost and accessibility are frequent barriers to use. Research into the effects of artificial light at night (ALAN) further compounds these difficulties with requirements for sensitivity to extremely low light levels across the ultraviolet to human-visible spectrum. Here I present a open-source spectroradiometry (OSpRad) system that meets the design challenges of typical ALAN research. The system utilises an affordable miniature spectrometer chip: the Hamamatsu C12880MA, and combines it with an automated shutter and cosine-corrector, microprocessor controller, and graphical user interface “app” that can be used with smartphones or desktop computers. The system is designed to be user-friendly, adaptable, and suitable for automation/data-logging. All code and 3D printed parts are made available open-source. I constructed 5 units and tested their linearity, spectral sensitivity, cosine-corrector performance, and low-light performance. There were modest unit-specific differences in spectral sensitivity, implying calibration is required for maximal accuracy. However, depending on the application, it may be acceptable to use a default calibration template together with careful experimental design considerations to mitigate unit-specific differences. All other performance characteristics were highly consistent. The OSpRad system was able to measure spectral irradiance down to around 0.005 lx, and spectral radiance down to 0.001 cd.m^-2^, meaning it would be able to measure night-time lighting under the vast majority of real-world conditions. The OSpRad system’s low cost and high sensitivity make it well suited to a range of spectrometry tasks in general, and ALAN research in particular.

The project is hosted on GitHub here: https://github.com/troscianko/OSpRad

## Introduction

The night-time light environment is characterised by extreme differences in spectral intensity and spectral composition compared to typical daytime lighting. This is due to temporal and spatial variation in natural light sources and artificial light at night (ALAN), together with highly variable degrees of atmospheric scattering and reflection. ALAN is a widespread and growing source of environmental pollution that is emitted by street-lights, homes and businesses, and it dramatically alters the timing and spectral composition of the night light environment (Gaston et al., 2013; Kyba et al., 2017; Sánchez de Miguel et al., 2022). The detrimental effects of ALAN have been demonstrated in a wide range of taxa and ecosystems (Owens & Lewis, 2018; Sanders et al., 2018), affecting animal behaviour (Becker et al., 2013), physiology (Dominoni et al., 2013), pollination (Knop et al., 2017; Macgregor et al., 2019), sexual signalling (Lewis et al., 2020), and population dynamics (van Grunsven et al., 2020). Measuring the spectral properties of the night-time light environment is therefore essential for ALAN research, however the technological requirements for this are typically beyond the abilities of standard spectroradiometric equipment, and cost is a limit to widespread data collection.

Animal visual systems vary substantially in their sensitivity to different ranges of spectra, and to different light intensities (Kelber et al., 2017). Like most terrestrial vertebrates, humans have retinas with cones that function in typical daytime light intensities, and rods that provide achromatic vision at low light levels. Even among species with these dual function retinas there is considerable variation in the degree of low-light sensitivity, with adaptations such as rod-dominanated retinas, powerful optics and reflective membranes that boost low-light sensitivity (Land & Nilsson, 2012). However, a growing number of species have been found to possess chromatic low-light vision, and sensitivity into the ultraviolet range (Kelber et al., 2017). This includes frogs and toads that have evolved cone-like-rods (Koskelainen et al., 1994; Yovanovich et al., 2017), geckos that possess three classes of rod-like-cones (Roth & Kelber, 2004), and nocturnal bees and hawkmoths that have the ability to discriminate colour down to star-light levels of illumination (Kelber et al., 2002; Warrant & Somanathan, 2022). Measuring the spectral composition of the night time light environment is therefore essential for understanding the potential effects on the vision of different species, and can be a useful tool for predicting the impact on various species’ visual ecology. For example, artificial light sources with narrow peaks in their spectral emissions were found to have unpredictable interactions between light intensity, object reflectance and colour discrimination for hawkmoths (Briolat et al., 2021). The spectral composition of light also affects a number of other key biological processes; for example shortwave light in particular is used by plants to detect photoperiod, and is used to regulate the circadian rhythms of many species (Bennie et al., 2016; Grubisic et al., 2019; Sánchez de Miguel et al., 2022). Taken together, this demonstrates the need for further research into the effects of ALAN using measurements that capture the ecologically relevant intensity, spectral, spatial and temporal properties of the light environment.

The nighttime light environment can be measured using a range of radiometric techniques that are reviewed by (Hänel et al., 2018), and I give a brief overview here with a focus on the principles likely to be important to biologists. 1-dimensional measurements such as those provided by lux meters are comparatively affordable and straightforward to acquire, yet they lack the spectral information likely to be important for the reasons outlined above. The spectral sensitivity of a lux meter is also often unknown, and unlikely to match the spectral response curves of humans or any other animal well. As such the measurements are unreliable when measuring light sources that have narrow peaks in their emission spectra (e.g. the exact location of that spectral emission peak relative to the device’s spectral sensitivity function will dramatically affect the measurement, either under- or over-estimating luminance). Low cost sensors are also typically unable to perform adequately under low-light conditions (e.g. below ∼10lx). Camera-based techniques are valuable because they provide both spatial and spectral information; when carefully calibrated, such systems can be used to quantify radiance across the entire visual scene (Nilsson & Smolka, n.d.), however the limited spectral range of camera systems means they are not well suited to accurate modelling of animal visual systems or other processes that are spectrum-specific. e.g. widely used calibrated photography methods that convert from camera sensitivities to animal cone catch require knowledge of the illuminant spectrum (Troscianko & Stevens, 2015), which is highly consistent in typical daytime, terrestrial conditions, but under most low-light conditions the illumination spectrum is often composed of unknown contributions of multiple artificial and natural sources. This can cause metameric effects whereby combinations of (potentially wildly) different illuminant spectra can combine with a single reflectance spectrum to create identical camera tristimulus (RGB) values. Therefore even if the camera’s spectral sensitivity functions are known it may be impossible to reliably or accurately infer cone catch values. Moreover, while basic camera-based systems are typically affordable, adapting them for ultraviolet radiometric measurements offers additional challenges and costs. Cameras converted to “full spectrum” operate across the UV, human-visible, and near infrared ranges. This is achieved by removing the camera’s own UV-IR blocking filter that sits in front of the sensor, and capturing repeat measurements through multispectral pass filters (e.g. visible-pass and UV-pass) (Troscianko & Stevens, 2015). Camera CMOS sensors are far less sensitive to UV than near-infrared ranges, therefore it is critical that UV pass filters effectively block all near IR spectra, and they typically have comparatively low transmission (the Baader Venus-U filter is typically used). Moreover, wide-angle lenses are not commercially available, presenting significant difficulties for whole-scene radiance measurements in UV and low-light. Spectroradiometry offers the highest spectral resolution of any method, and can be used to either measure spectral radiance or spectral irradiance (with the use of a cosine corrector). Spectroradiometers allow spectral resolution to be sacrificed for better low-light sensitivity by using larger aperture slits, so systems are typically configured for the anticipated use (high sensitivity or high spectral resolution). Although radiance is only measured in a narrow incident angle, mechanical controllers can be used to build up spatial information across the night sky (Hänel et al., 2018) or build up a hyper-spectral image. While spectroradiometry is generally considered the gold standard for radiometric measurements, their widespread use by biologists remains limited, and this is particularly true for those wishing to measure ALAN or natural low-light levels across the UV and human-visible portions of the spectrum. The availability and cost of commercial systems that meet the typical requirements of ALAN researchers acts as a real impediment to progress in the field. This is particularly true for data collection over large spatial and temporal scales, and in low-income nations where the effects of ALAN could present significant yet unknown risks to biodiversity.

The Hamamatsu C12880MA is a micro spectrometer that is principally designed for integration into medical components. However, it’s specifications appear well suited for use in biological and ALAN research with a spectral range of ∼320 to 850nm, which covers the wavelengths relevant to key biological processes. It is also designed to offer good low light sensitivity with a 288-site CMOS sensor, integrated optics and comparatively large 500 micron slit size. It’s typical spectral resolution according to the manufacturer (full width at half maximum, FWHM) of ∼9nm (maximum of 15nm) is also well suited to biological applications where a higher spectral resolution is rarely necessary given typical receptor spectral sensitivity curves. The unit is commercially available at low cost (around £220 at the point of writing; Hamamatsu UK). However it is only available as a bare electrical component that requires considerable further integration with circuitry that can control the sensor and read the output (off-the-shelf prototyping boards are considerably more expensive than the units themselves). Moreover, in its basic state it is not well suited to automated low-light measurements that require controlling for the sensor’s non-linearities and dark response.

In this paper I describe and test an open-source spectroradiometry (“OSpRad”) solution for researchers that can meet the design challenges described above at an exceptionally low-cost. This utilises 3D printed parts, an open source microcontroller, automated radiance and irradiance measurement capability (utilising a cosine corrector and shutter), and a user interface capable of running the device from a low-cost smart phone or almost any computer capable of running Python and connecting via USB. The interface also provides a straightforward repeat timer function for automated data logging, and can easily be modified and integrated with other equipment. First I describe the design and construction, then I calibrate and test the performance of multiple OSpRad devices.

## Design

### Spectrometer Controller

The C12880MA chip must be interfaced through a series of input/output electrical pins that control the integration time (the time over which light is collected) and provide output from each of the 288 photosites. Each photosite’s peak wavelength sensitivity is described by a function unique to each device, provided by the manufacturer. CMOS photosites increase their voltage linearly with the energy of light they receive; this voltage is then converted to a digital count via an analogue-to-digital-converter (ADC). I used an Arduino Nano microcontroller to interface with the C12880MA because they are small, affordable and readily available (the Nano units used here were manufactured by Elegoo, or unbranded manufacturers). The OSpRad unit receives power and communication via the Arduino’s USB port, while a serial connection to a desktop computer or smartphone is used to send commands and receive data. I developed the code (“firmware”) for the Arduino based on the manufacturer’s specifications, partially inspired by existing open-source code (*GitHub - Impfs/Review*, n.d.). The ADC on the Arduino Nano offers 1,024 linear levels, and the sensor’s output voltage range limits this to approximately 900 linear levels. This would give the system a maximum theoretical dynamic range of 900:1, which can be improved dramatically by averaging over multiple exposures, and will also be reduced under noisy low-light situations. The controller’s default behaviour is to record up to a maximum of 50 exposures, which it then averages at each photosite for each measurement. When 50 exposures would cause the measurement to take longer than 1 second, the code reduces the number of averaged exposures to be less than one second in total, down to a minimum of 3 exposures. The firmware automatically estimates the ideal integration time by incrementally taking test exposures and doubling the integration time until any part of the spectrum reaches the saturation point. Once saturation is reached, the firmware uses the preceding (unsaturated) measurement to calculate the exposure that would cause the peak intensity measurement to be 0.8 of that which would cause saturation. The maximum exposure is 60 seconds, above which the system would need additional stability testing (initial testing suggested some photosites may suffer voltage collapse). As with any CMOS sensor, longer exposures increase the noise associated with measurement, and with low light levels the voltage at each site that represents “zero” (the black point) is uncertain. Therefore, following each measurement (which will be an average of multiple exposures as described above), the firmware closes a mechanical shutter and takes a dark measurement using an identical exposure regime (the same integration times and numbers of exposures), and uses this to calculate the black point at each site. ADC counts *c* at each photosite *p*, are calculated as:

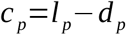

where *l* is the light measurement and *d* is the dark measurement. This controls for temperature and voltage fluctuations that affect the sensor’s black point. When exposures are longer than 1 second the dark measurements are interleaved between light measurements, so that long exposures have closer temporal matching of black point measurements. In addition to this automated exposure behaviour, the user can also manually control the integration time, number of scans, and frequency of repeat measures (and whether repeat measures are radiance, irradiance, or both).

### Electronics & Housing

The housing was created in Blender (version 1.92), and was designed to make device small, robust, easy to print, and with the potential for waterproofing (for data logging, see below). I recommend printing in black PLA or ABS, and ensuring the shutter blocks near infrared light (e.g. black PET often transmits near IR). The shutter is controlled by a digital servo (Savox SH-0256), which was chosen because of its low-cost, availability and relative simplicity. Each servo has a slightly different positional response to the pulse-width-modulated position signal from the Arduino nano. Each OSpRad unit therefore requires the servo’s three shutter wheel positions (closed, radiance, irradiance) to be calibrated when first constructed.

## Construction

Construction of this device requires some soldering, access to 3D printed parts (figure 1, there are services that offer this through the internet), and a computer that can run the Arduino IDE software (Linux/Windows/MacOS) to write the firmware code to the Arduino nano. Please consult the construction guide for detailed assembly information. The housing was designed to be compact with the potential to make it weatherproof/waterproof for remote logging installations. Waterproofing could be achieved either by painting the 3D printed housing, or printing with ABS plastic and using an acetone vapour bath to fuse the surface. Rubber O-rings are also required for waterproofing the seals (which could be purchased, or made from a sheet of silicone) The user should must also select the desired level of protection for the aperture. For purely lab-based measurements the unit will not require any physical cover for the aperture, and using no cover will improve the performance of the cosine-corrector at shallow angles (see figure 6). However, I chose to use protective plastic covers (circles of 16mm diameter, 2mm thick, made from UV-transmitting sunbed-grade PMMA, Bay Plastics Ltd.), cut with a laser cutter. The material can easily be cut into square tiles that would also be suitable. Fused silica microscope cover slips would also function well as protective covers, and would have slightly superior transmission (though would be less impact resistant). The cosine corrector can also be made in different ways. An ideal cosine corrector simply scatters transmitted light in all directions equally, resulting in illuminance of it’s surface facing the spectrometer that follows a cosine-function. I made units that either had a disk of 0.5mm thick virgin PTFE (Bay Plastics Ltd.), sanded on both sides with 180 grit sand paper in a circular motion, or by sandwiching 4 layers of plumber’s PTFE tape (Silverline) in different directions at each layer.

**Figure 1.**
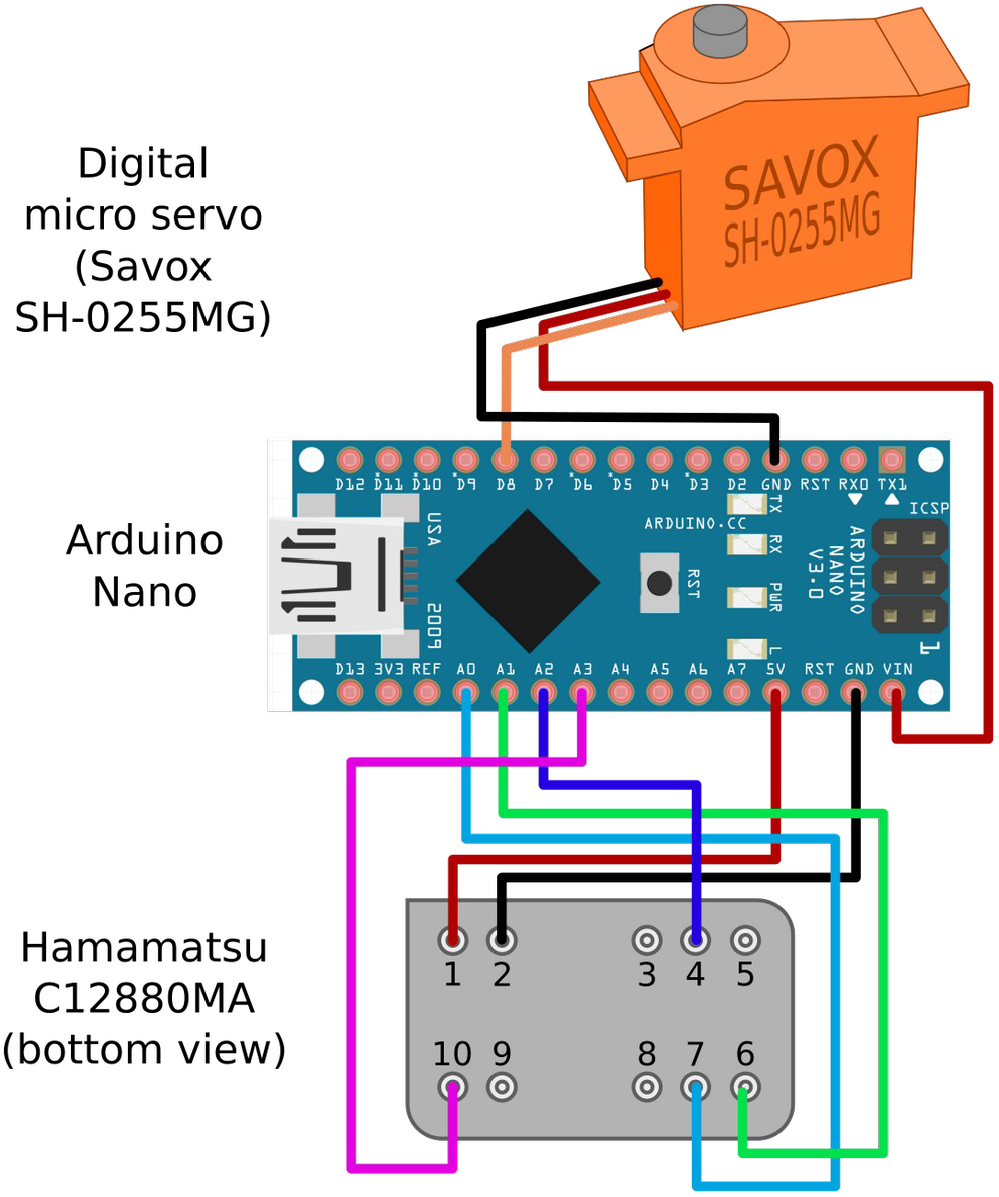
Circuit diagram, showing connections between the microcontroller (Arduino Nano), spectrometer chip (C12880MA) and digital servo that controls the shutter/filter wheel. Note that the C12880MA chip and servo run from separate voltage rails, due to the voltage drop in the VIN’s diode.

Further optional equipment includes a calibrated light source for more accurate absolute measurements (see below), and an Android smart phone or computer to connect to the spectroradiometer via USB serial interface.

## User Interface

I developed a graphical user interface application (“app”) that can control OSpRad units from either a desktop computer or a smartphone, making the system highly portable, and flexible. The app was designed to be easy to use, and suited to both point measures and regular data logging. When a user clicks either “Radiance” or “Irradiance” buttons, the app uses the default automated measurement behaviours described above. However the user also has options for manually controlling the integration time and number of scans to average. Regular repeat measures can be made automatically by ticking the check-box, with a field for inputting the time delay between measurements (the default shown in figure 3 is 300 seconds). Further check-boxes allow the user to specify whether repeat measures should be radiance, irradiance or both. A graph shows the spectrum of the previous measurement (Le or Ee values), together with calculations of luminance (in cd.m^-2^) or illuminance (lux, lm.m^-1^) based on the CIE 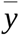 2006 analytical function (the app generates the shape of the Y function to match each unit’s spectral response). The number of saturated photosites are also shown in the “Sat.” field. Any photosites with saturated values will give under-estimates of the flux, and importantly, saturation can only be determined from the light (*l*) measures above (before the dark values are subtracted). As such, any measurements that have multiple saturated values should be treated cautiously and avoided where possible. Clicking the “Save” button will save the current measurement, appending it to the data file. After saving, the app shows a list of recently saved spectra, and prevents re-saving until a new measurement is made.

**Figure 2.**
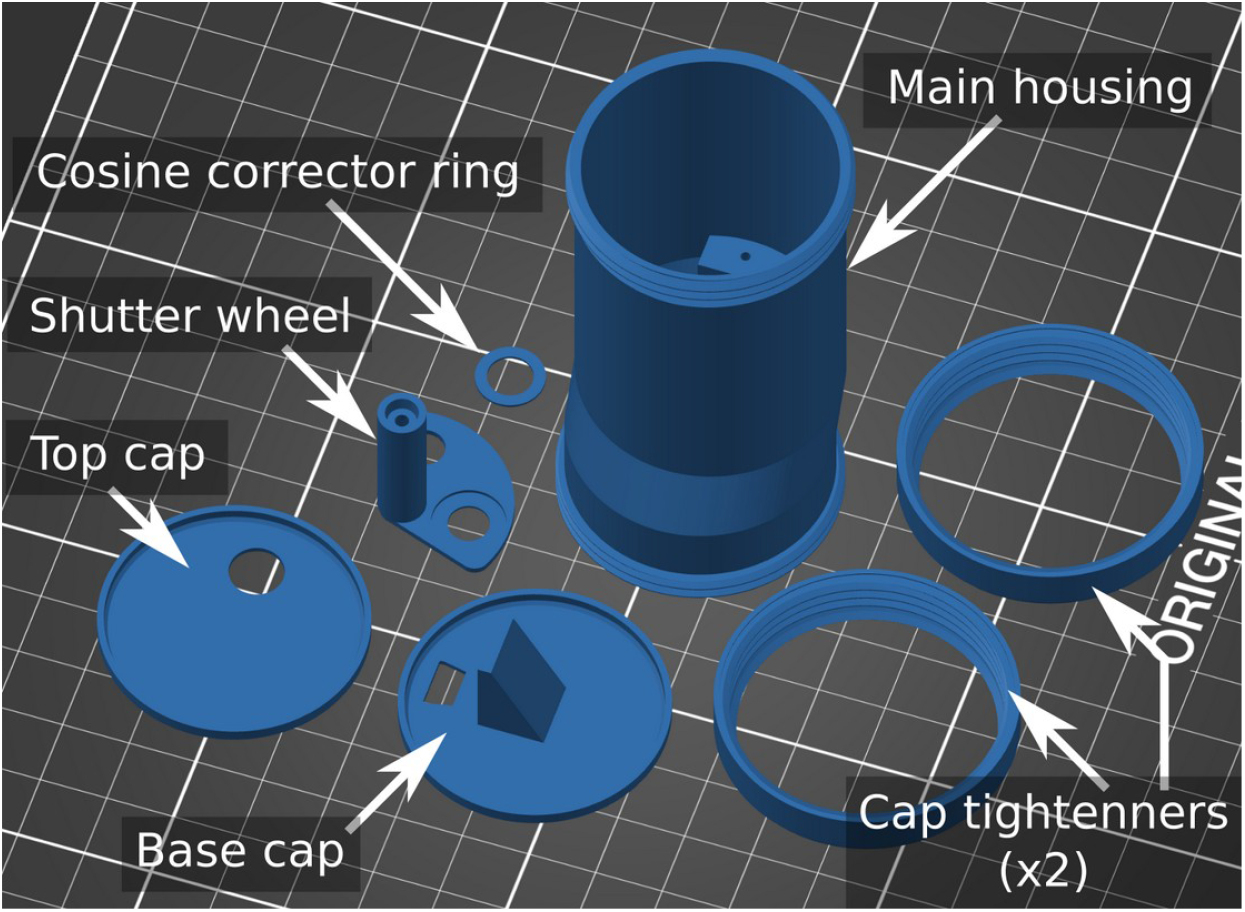
3D printed components required for construction.

**Figure 3.**
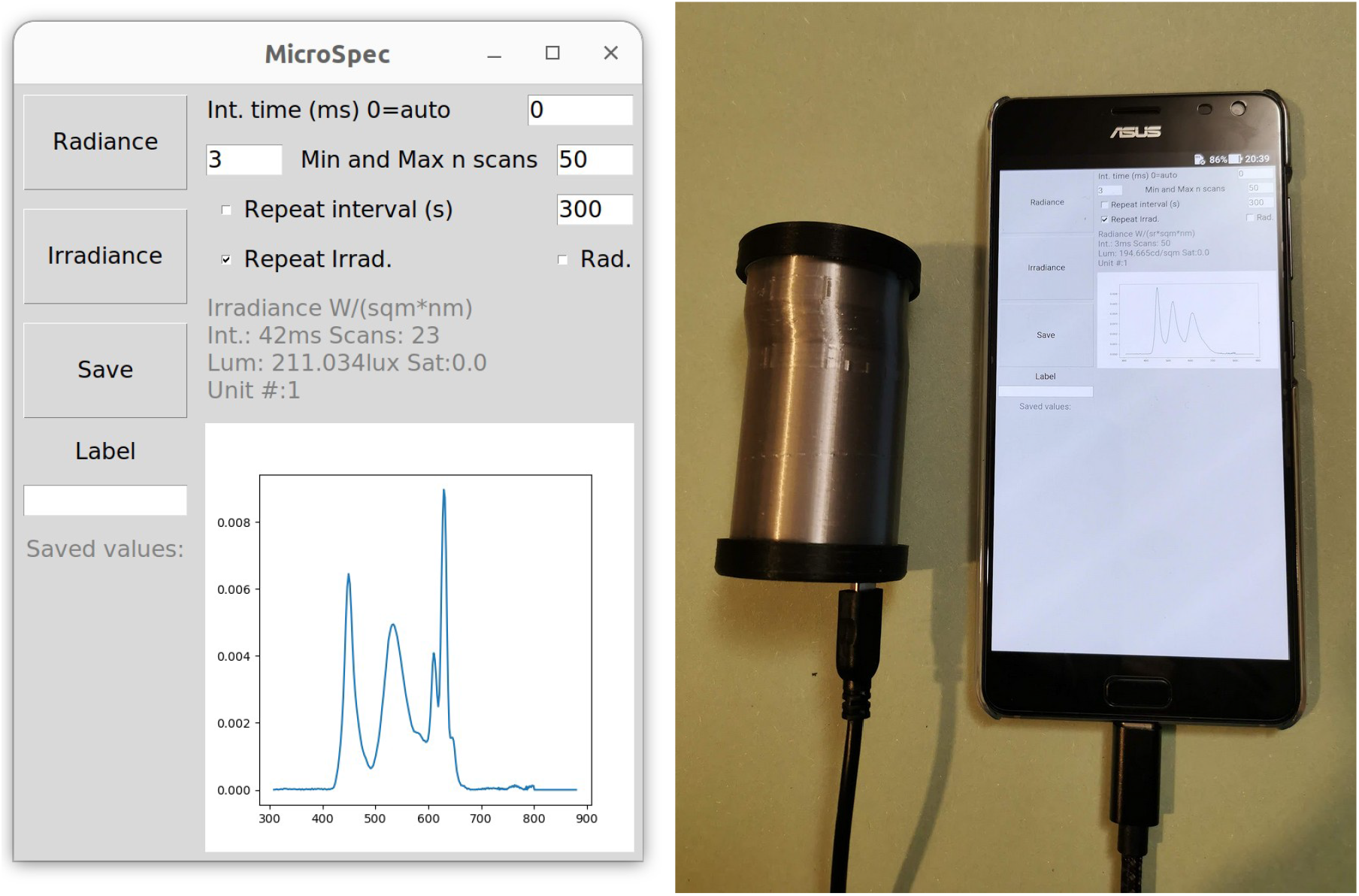
App with graphical user interface for controlling OSpRad units. Left shows the software running on a Linux desktop, and right shows the same code running on an Android smart phone (connected via a standard USB-c to female USB type A cable).

Data are saved in a manner designed to be both user-friendly and non-destructive. Along with calibrated spectral data (Le [W.sr^-1^.m^-2^.nm^-1^] or Ee [W.m^-2^.nm^-1^] values at each photosite), and wavelength data for each photosite, the app also saves the unit identifier code (which is used for looking up unit-specific calibration data), the user-specified label, the time and date of the recording, the type of measurement (radiance or irradiance), the integration time and number of scans, the number of saturated values, and also the raw spectral count data (*c*). These raw count data have not been calibrated, meaning any measurement can be re-calibrated post-recording. This is potentially useful if a researcher needs to take spectral recordings before they are able to fully calibrate the equipment, or to apply a dark correction (see below). The data are all appended to a “data.csv” file in the same directory as the app’s code. The app also requires calibration data to be present in the same directory, stored in a file called “calibration_data.csv”. This contains the wavelength, linearisation and radiance/irradiance spectral sensitivity calibration data and coefficients for each unit.

The app was developed in IDLE (version 3.10.6), and written in *Python* (version 3.10.6), with the graphical user interface based on *tkinter* (version 8.6.12). The code has a limited number of additional dependencies that need to be installed alongside Python 3:, *matplotlib, usbserial*, and (for Android only) *usbserial4a*. On Android smartphones the code can be run via the free Pydroid 3 App (see figure 3). Adapting the code to support Windows or MacOS will also be straightforward by adding system-specific serial connection details.

## Results

### Calibration

Measurements of absolute radiance or irradiance require two main types of calibration; linearisation (ensuring sensor responses scale linearly with flux), and spectral sensitivity (which is a function of optical transmission, diffraction grating efficiency, and CMOS sensor sensitivity). This calibration should only be performed once the entire system has been assembled (including the cosine corrector and any protective cover).

#### Linearisation

There was significant deviation from linearity at lower pixel counts (lower counts under-estimated the flux), which needs to be controlled for to ensure low-light measurements do no under-estimate low-light intensity. Linearity was modelled by measuring the radiance of an AMOLED screen (smartphone, ASUS A002), placed directly against the OSpRad’s protective cover. Measurements were taken over multiple integration times in one-octave steps, ramped down from saturation point (typically ∼100ms), to 1ms (minimum value), and then back up to saturation point (ramping both ways to ensure the light source remained stable). Each unit’s summed counts, 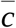, across the spectral emission range of the light source were calculated as:

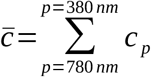

where *c* is the ADC response (count) at each photosite *p*, for sites between 380 and 780nm (the approximate spectral emission range of the source). The expected linear count rate 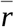 is calculated from exposure time *t*:

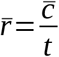

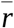 is then scaled to max=1 for comparison between units. 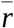 was found to show a log-linear response to 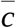, which could be modelled accurately with the function:

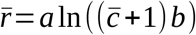

where *a* and *b* are coefficients fitted with least-squares regression, shown in table 2. Fitting was performed in ImageJ (version 1.53). This function levels out as counts approach zero, which reduces noise compared to typical linearisation functions such as the Gamma variate, which also fits the linearisation function well, but causes considerable artificial noise as counts approach zero. Linearised counts at each photosite are then calculated as:

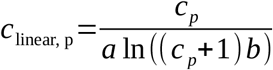

**Table 1.** List of parts.

| Part | Count/<br>amount | Typical net<br>cost (£) |
| --- | --- | --- |
| Hamamatsu C12880MA | 1 | 220 |
| 3D printed components (black PLA or<br>ABS recommended, not PET due to<br>IR transparency) | 1 | 1 |
| Arduino Nano | 1 | 5 |
| Diffuser: 0.5mm sanded virgin PTFE<br>sheet (recommended)<br>or plumber’s PTFE tape | 8mm disk<br>or 1 roll | 5 |
| Digital micro servo (Savox – SH-0256<br>recommended) | 1 | 15 |
| USB cable(s) for mobile phone. e.g.<br>USB-C to USB-B female, and USB-B<br>male to USB-mini | 1 | 10 |
| Solder and cables | - | - |
| UV curing glue | <5ml | 5 |
| Circular fused silica microscope<br>cover-slip or UV-transmitting PMMA<br>disk (optional) | 1 | 5-25 |

**Table 2.** Linearisation coefficients and model fits for five OSpRad units, fits are also shown in figure 4.

| Unit | $a$ | $b$ | $R^2$ |
| --- | --- | --- | --- |
| A | 0.12828 | 2.16327 | 0.9891 |
| B | 0.12249 | 3.258 | 0.9913 |
| C | 0.14231 | 1.06125 | 0.9984 |
| D | 0.13812 | 1.37537 | 0.9959 |
| E | 0.12032 | 3.89444 | 0.9976 |

Any negative count values are assumed to follow the same function (the sign is switched, and switched back following linearisation).

#### Spectral Sensitivity

Spectral calibration should be performed using a light source with a known/measured emission spectrum. I attempted this with a Deuterium source (Thorlabs SLS204), however, the large 650nm spike reduced the effective dynamic range in order to avoid saturation. I also used a halogen source, however, this was insufficient for calibrating the lower UV range and its extreme near infrared output appeared to cause second order diffraction grating interference. I therefore used a standard xenon photography strobe flash (Neewer NW-14EXM) that had been converted to full-spectrum by removing the plastic (UV-absorbing) protective covers and replacing them with diffuse 0.5mm virgin PTFE sheet. The xenon source was used to illuminate a virgin PTFE sheet (20×20cm, 1mm thick) from behind to create a uniformly illuminated surface in a dark room. Radiance and irradiance were measured with a Jeti Specbos (1211UV) spectroradiometer with NIST-traceable calibration, ∼20cm from the rear-illuminated surface. The OSpRad units were then used to measure the radiance and irradiance of the same surface from the nearest distance that did not cause saturation (∼100cm away; this light source is strobed, so integration time cannot be used to limit exposure and the C12880MA chips have far higher sensitivity than the Specbos). This process allowed for calibration of the relative spectral sensitivity of the units, but given the differences in detection distance, could not give absolute values. Absolute radiometric calibration was next performed using an AMOLED screen (as above, ASUS A002). The Specbos and OSpRad units were placed directly against the screen, which was displaying a white surface, for measurements of absolute intensity. Calculations for all these measurements are included on GitHub (see below). Following calibration, radiance Le [W.sr^-1^.m^-2^.nm^-1^] is calculated at each photosite’s peak wavelength (λ) as:

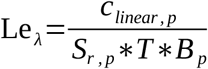

where *c*_*linear, p*_ is the average linearised count at each photosite *p*; *S*_*r*_ is radiance sensitivity; *T* is integration time (milliseconds) and *B* is the spectral width of the bin (nanometres). Irradiance Ee [W.m^-2^.nm^-1^] is calculated similarly, substituting *S*_*r*_ for *S*_*i*_ (irradiance sensitivity). The sensitivity functions, smoothed using a Gaussian filter (σ = 3) are shown in figure 5. Note that these units are watts per nanometre, so to recover the energy of a given spectrum at each photosite, Le or Ee must be multiplied by the bin width *B* at that photosite. Wavelength λ is calculated from *p* using a function with coefficients supplied by the manufacturer for each unit.

**Figure 4.**
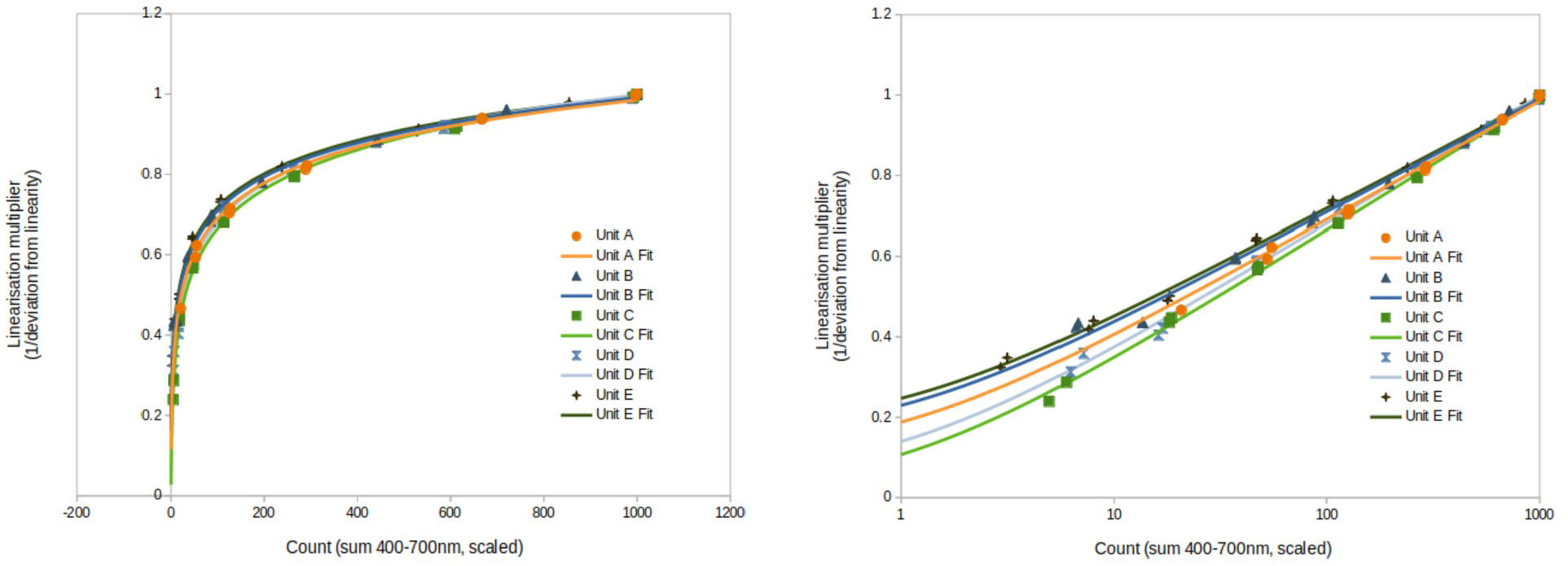
Linearisation modelling of five OSpRad units shown with linear (left) and log (right) x-axes. X-axes show the sum of counts 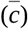 from 380-780nm, of a stable light source measured at a range of integration times, then scaled to max=1000 (the saturation point at each pixel). The y-axis shows the linearisation multiplier 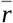 (count values are linearised by dividing them by this value, see text).

**Figure 5.**
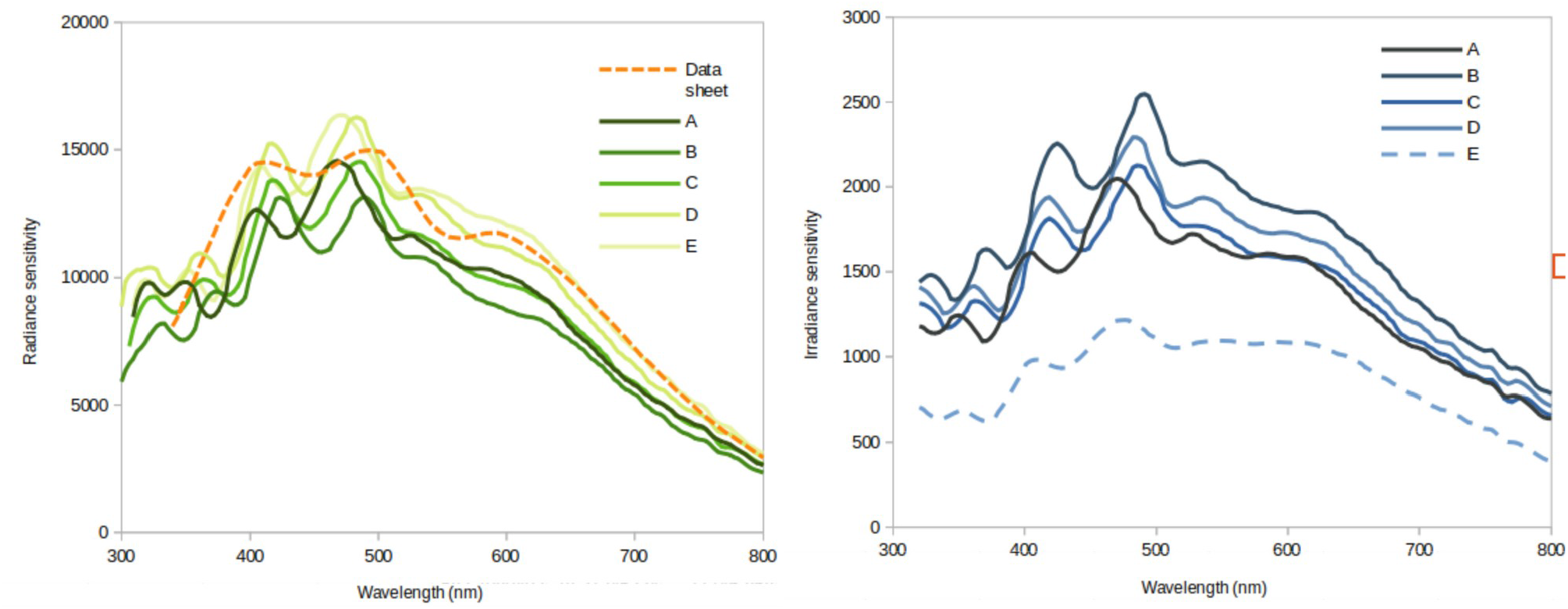
Radiance (left, *Sr*) and irradiance (right, *Si*) sensitivity functions for five OSpRad units. The orange dashed line shows the sensitivity published by the manufacturer (scaled to match the approximate sensitivity y-axis of the plot). Note that the lower irradiance sensitivity of unit “E” is due to a difference in its cosine-corrector construction (4 layers of PTFE tape, where units A-D use a single 0.5mm PTFE sheet sanded with 180 grit paper).

#### Cosine corrector

The cosine correctors were tested by illuminating each unit with a stable LED point light source (Phillips PC amber Luxeon) with a continuous current of 152mA (∼2.7v). The LED was attached to an arm that rotated around the OSpRad’s cosine corrector with a radius of 660mm. The OSpRad’s surface was levelled with a spirit-level, and the arm’s angle was measured with an inclinometer. Irradiance was measured from angles of 0 (directly above the surface) to 80 degrees and then back up to 0 degrees, in 10 degree increments. Default measurement behaviour was used (auto-exposure, minimum of 3 scans). Irradiance was measured with the plastic protective covers removed to test optimal performance, and then again with the plastic covers in place. Results are shown in figure 6. The bare 0.5mm PTFE cosine corrector performs close to the ideal cosine function down to roughly 70-80 degrees, below which the housing casts a shadow over the surface. The PFTE tape-based cosine corrector has slightly poorer performance. Performance is reduced by the presence of the plastic cover in all cases (due to specular reflection and refraction at acute angles, particularly below 50 degrees). Nevertheless, performance is highly consistent among all four 0.5mm PTFE cosine correctors.

**Figure 6.**
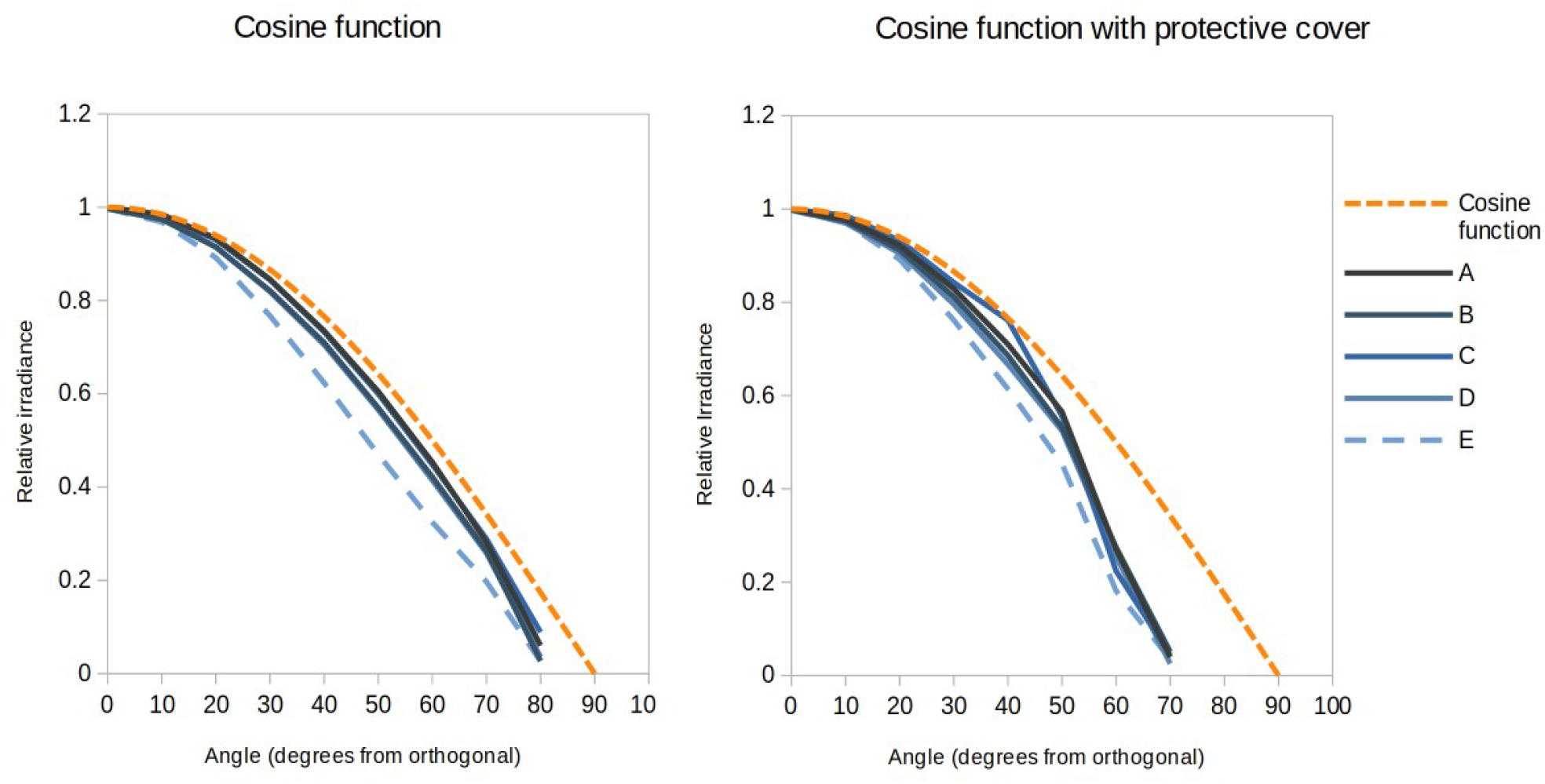
Cosine corrector test data, with a bare surface (left) or plastic protective cover (right). An ideal cosine corrector would cause irradiance to fall off with the angle of illumination following the cosine function (shown in dashed orange). OSpRad units A-D use 0.5mm thick sanded PTFE filters, while E uses four layers of PTFE tape.

#### Low light performance

Low light performance was tested with an AMOLED screen (ASUS A002, as above), displaying a white square on a black surround, measured from 100mm away. Screen brightness was reduced to its minimum, and the size of the square was reduced incrementally to assess the lowest light levels that produced usable spectral data. The three clearly defined emission peaks of the screen’s LEDs are shown in figure 7. The level of acceptable noise will depend on the intended use of the resulting spectra, and can generally be reduced by averaging multiple exposures, and must ultimately be assessed by researchers. As such I present unfiltered raw count data together with processed spectra for assessing low-light performance.

**Figure 7.**
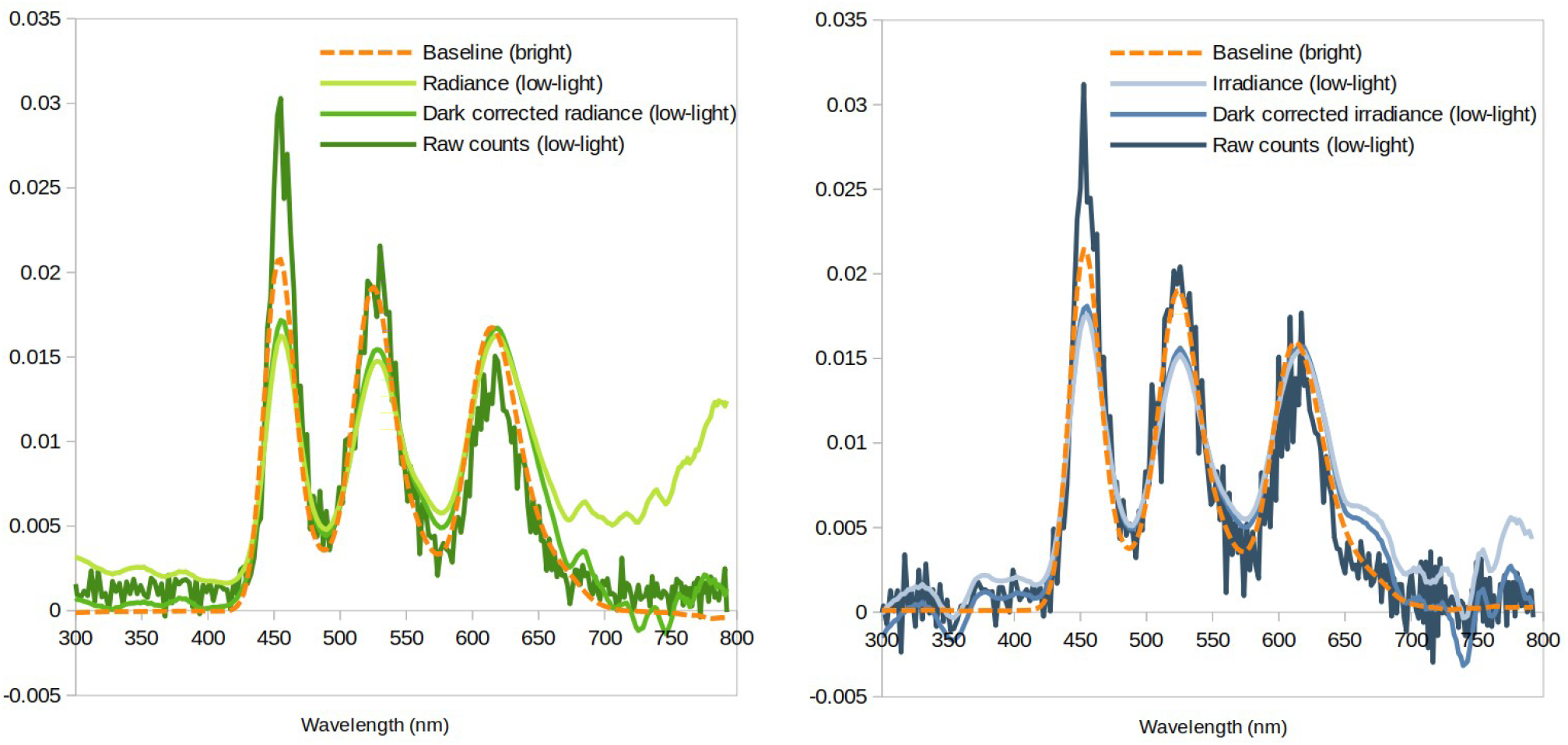
Low-light performance testing for radiance (left) and irradiance (right), showing the baseline measurements (taken under bright light conditions of 5.48 cd.m^-2^ and 49.1 lx respectively), raw counts 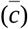, calibrated radiance or irradiance values, and dark-corrected values. Radiance in this example was 0.0013 cd.m^-2^ (measured by averaging 80 scans of 30 seconds), and irradiance was 0.0051 lx (average of 48 scans of 30 seconds). All metrics are scaled to area-under-curve from 425-650nm = 1 for assessing relative spectral shape. All spectra shown were smoothed with a Gaussian function (sigma=3), with the exception of raw counts. Note that raw counts do not account for spectral sensitivity or linearisation.

Irradiance measurements use the cosine corrector, which reduces sensitivity to low-light. As such, radiance measurements will allow for lower-light performance than irradiance measurements. This testing used an OSpRad unit with a 0.5mm sanded PTFE cosine corrector (unit “B”). Usable radiance spectra were measured down to 0.0013 cd.m^-2^, and irradiance spectra were measured down to 0.0051 lx (see figure 7). Under these extreme low-light conditions there was an increase in estimated near-infrared energy caused by an under-estimate of the dark value and subsequent amplification of noise in the low-spectral-sensitivity regions of the spectrum (infrared and UV). This is likely caused by scattered light inside the spectrometer, so cannot be accounted for by closing the shutter. To compensate for this effect, a dark correction was applied by subtracting 0.5 from raw radiance counts, and 0.2 from raw irradiance counts in the examples shown in figure 7.

## Discussion

OSpRad spectroradiometers based on Hamamatsu C12880MA chips offer a cost-effective solution for spectral radiance and irradiance measurements in the UV-A to near infrared range at low light levels. This makes them particularly well suited to behavioural and ecological measurements, and investigating the effects of ALAN.

Low-light sensitivity of the OSpRad units was found to be around 0.005 lx for irradiance, which is roughly between starlight and the lowest levels of moonlight (Hänel et al., 2018; Johnsen et al., 2006). Sensitivity was higher for radiance, measuring spectra at around 0.001 cd.m^-2^, meaning it is able to measure spectra under the vast majority of typical night-time light intensities; for example, the recommended luminance of street lights is 0.3-2 cd.m^-2^, and the milky way is ∼0.0027 cd.m^-2^ (Hänel et al., 2018). Nevertheless, further work could assess whether low-light sensitivity can be improved through hardware or software solutions.

Spectral calibration of OSpRad units is a process that requires access to specialist, and expensive equipment that could be a barrier to their widespread use. The data presented here from 5 units suggests that absolute sensitivity is comparatively uniform among units, whereas there are modest differences in spectral sensitivities, particularly below ∼500nm. Therefore, if researchers require high levels of confidence in the accuracy of their recordings, each unit should be calibrated independently. However, if researchers were to use the data presented here for their otherwise uncalibrated units, the overall luminance/illuminance measurements produced would be comparatively accurate (particularly when considering the log-scale of light intensities that they will typically be used to measure). Exact measures of spectral intensity below ∼500nm would be prone to error, however the degree of error will be highly dependent on the shape of the spectra being measured and spectral sensitivities of the visual system. Careful experimental design may also be used to rule out major systemic bias as a result of imperfect unit-specific calibration. For example, an uncalibrated OSpRad units could be used to reliably measure relative spectral shifts (e.g. proportions of UV to SW light) with high repeatability. The use of uncalibrated equipment is most problematic if colour measurements are being compared among multiple units with slight (unknown) differences in their spectral sensitivity. One possible solution to this would be to calibrate all OSpRad units against a single “master” unit and accept some degree of absolute error, while maintaining cross-unit repeatability. Additionally, the spectral sensitivity functions (figure 5) appear to vary more in wavelength offset than overall shape, implying unit-specific calibration could be achieved by fitting the curves presented here to measurements of a spectrally consistent source (such as a halogen bulb), shifting a template sensitivity curve along until peaks are eliminated. Linearisation fitting shows that all units are behaving in a similar manner, with modest differences between units.

Cosine correctors that match a true cosine scattering performance are typically extremely expensive components constructed from materials such as Spectralon sintered PTFE. For the OSpRad I sought alternatives that could readily be used in low-cost or data-logging applications, where perfect cosine performance is typically not critical. Nevertheless, I demonstrated that a cosine corrector made from sanded 0.5mm thick virgin PTFE sheet performed well in both transmission and scattering of light. The cosine corrector created from four layers of PTFE tape resulted in slightly poorer transmission and diffusion characteristics.

Future developments could investigate performance improvements such as high-light-intensity performance (sub-millisecond integration times will be required to measure sources >∼700 cd.m^-2^), high dynamic range measurements, or use of a mechanical gimbal to generate hyperspectral images/spatial information. In releasing these open-source tools I hope to facilitate research into monitoring and mitigating for the effects of artificial light at night.

## Source code and data availability

The project is hosted on GitHub here: https://github.com/troscianko/OSpRad and is released under a GPL-3.0 license. Hosted on GitHub are the 3D printed part designs, Arduino code, Python interface app, and the calibration data and calculations used in this paper. The initial release is also available here: DOI: 10.5281/zenodo.7419032

